# Two-photon calcium imaging of medial prefrontal cortex and hippocampus without cortical invasion

**DOI:** 10.1101/119404

**Authors:** Masashi Kondo, Kenta Kobayashi, Masamichi Ohkura, Junichi Nakai, Masanori Matsuzaki

## Abstract

*In vivo* two-photon calcium imaging currently allows us to observe the activity of multiple neurons up to ∼900 μm below the cortical surface without cortical invasion. However, many other important brain areas are located deeper than this. Here, we used a 1100 nm laser, which underfilled the back aperture of the objective, and red genetically encoded calcium indicators to establish two-photon calcium imaging of the intact mouse brain and detect neural activity up to 1200 μm from the cortical surface. This imaging was obtained from the medial prefrontal cortex (the prelimbic area) and the hippocampal CA1 region. We found that the neural activity related to reward prediction is higher in the prelimbic area than in layer 2/3 of the secondary motor area, while it is negligible in the hippocampal CA1 region. Reducing the invasiveness of imaging is an important strategy to reveal the brain processes active in cognition and memory.

## Introduction

Two-photon calcium imaging reveals the *in vivo* activity of multiple neurons at a cellular and subcellular resolution (Jia et al., 2010; Ohki et al., 2005). Recent work demonstrated that, by exciting red-fluorescent calcium indicators with a laser at wavelengths of 1000–1100 nm, it is possible to image neural activity in the mouse sensory cortex at depths of 800–900 μm from the cortical surface attached to a cranial window (corresponding to layers 5 and 6) (Dana et al., 2016; Tischbirek et al., 2015). However, for functional imaging of deeper regions, such as the medial prefrontal cortex, hippocampus, and basal ganglia, it has been reported that invasive penetration is unavoidable; it is necessary to insert a microlens or a microprism into the cortical tissue, or remove the cortical tissue above the target region (Attardo et al., 2015; Dombeck et al., 2010; Low et al., 2014; Pilz et al., 2016). In this study, we demonstrate that, by shortening the light-path length within the tissue to reduce light scattering (Helmchen and Denk, 2005) and exciting red-fluorescent genetically encoded calcium indicators (red GECIs; Dana et al., 2016; Ohkura et al., 2015), we could detect the activity of multiple neurons in the medial prefrontal cortex (the prelimbic [PL] area) and hippocampal CA1 region at depths of 1.0–1.2 mm in behaving mice, without the need for invasive penetration or removal of cortical tissue.

## Results and Discussion

The light-path length within the tissue was shortened by reducing large-angle rays emitting from an objective with high numerical aperture (1.05 or 1.00). To do so, the back aperture of the objective was underfilled with a diameter-narrowed laser beam (7.2 mm, in comparison with that of the back aperture of 15.1 mm or 14.4 mm) (Helmchen and Denk, 2005; Matsuzaki et al., 2008) at a wavelength of 1100 nm. Three to four weeks after an injection of adeno-associated virus (AAV) carrying the R-CaMP1.07 gene into the intact medial frontal cortex (mFrC) of 2–3-month-old mice, we observed R-CaMP1.07-expressing neurons in the mFrC at depths of 100–1200 μm from the cortical surface, in awake and head-restrained mice (Figure 1a, b and Video 1). Using a laser intensity of 170–180 mW under the objective, we could detect calcium transients at depths of 1.0–1.2 mm (Figure 1c). Cell death, severe inflammation, and heating-induced responses were not apparent on histological staining after long-term imaging of the mFrC (Figure 1d–g). This is consistent with a study that reported that heat-induced cell responses occur when the laser intensity exceeds 300 mW (Podgorski and Ranganathan, 2016).

**Figure 1.**
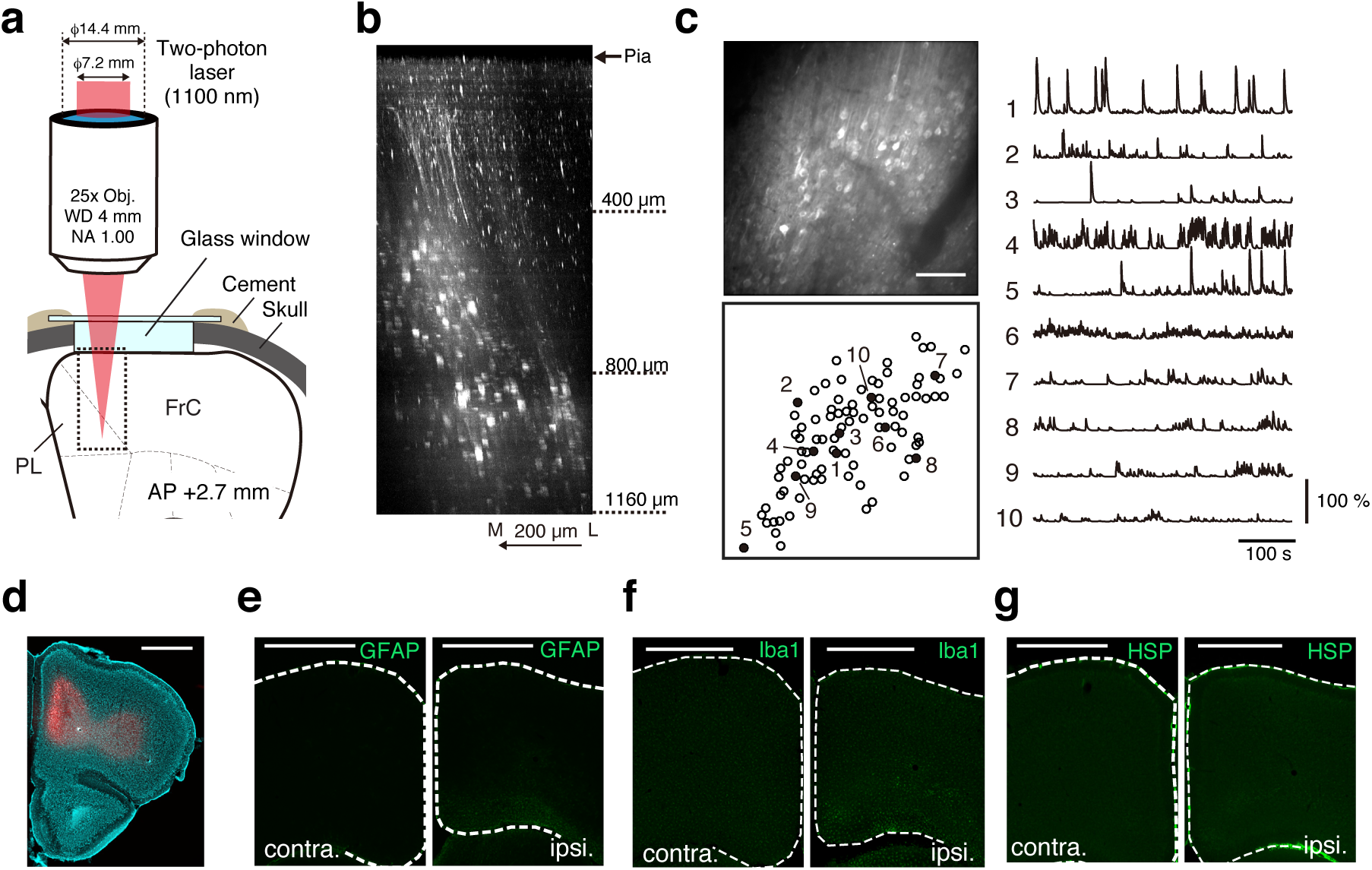
Optical access to intact medial frontal cortex. (**a**) Schematic illustration of *in vivo* two-photon imaging of the mFrC. The dotted square indicates the location where the Z-stack images, as in (**b**), were acquired. FrC, frontal cortex; PL, prelimbic area. (**b**) Representative XZ image of mFrC expressing R-CaMP1.07. Scale bar, 200 μm. (**c**) Left, top: representative time-averaged XY image of mFrC expressing R-CaMP1.07 at a depth of 1030 μm. Scale bar, 100 μm. Left, bottom: spatial distribution of identified neurons. Traces of the calcium transients of the numbered filled circles are shown on the right. (**d**) Expression of R-CaMP1.07 (red) and cell nuclei (NeuroTrace 435, cyan) after imaging of 11 fields at depths of 340–1140 μm for 5 days. The total duration of imaging was 200 min. There is no apparent abnormality of fluorescence expression in the mFrC. Scale bar, 1 mm. (**e–g**) Expressions of GFAP (**e**), Iba1 (**f**), and HSP70/72 (**g**) in contra-lateral (left) and imaged (right) hemispheres from the same mouse imaged in (**d**). Only slight glial activation and heat shock reactivity was observed. Scale bar, 1 mm.

We next examined whether neural activity in the hippocampus can be imaged without removal of the neocortical tissue lying above it (Figure 2a). CA1 GFP-expressing neurons can be detected by two-photon microscopy in 4-week-old mice, but not in 6- to 9-week-old mice (Kawakami et al., 2013). We therefore injected AAV-jRGECO1a into the hippocampus of mice aged between 12 and 14 days, and then performed imaging after another 2 weeks. When we deepened the focal plane below the white matter to depths of 900–1000 μm, we observed densely distributed fluorescent neurons typically located in CA1 pyramidal layer (Figure 2b, c and Video 2), as described previously (Dombeck et al., 2010), and clearly detected spontaneous calcium transients from these neurons (Figure 2c). No cell death or strong damage was apparent after 15 min of imaging (Figure 2d, e). By contrast, we could not detect any neural activity in the CA1 region of the mice when they were 3 months old. To image neural activity in the hippocampus or the infralimbic area of adult mice, it is necessary to use higher average power of the laser or higher peak power per pulse than that used in this study (Kawakami et al., 2015). In addition, adaptive optics (Ji et al., 2010) and further improvement of red GECIs (Dana et al., 2016; Inoue et al., 2014) will certainly be helpful.

**Figure 2.**
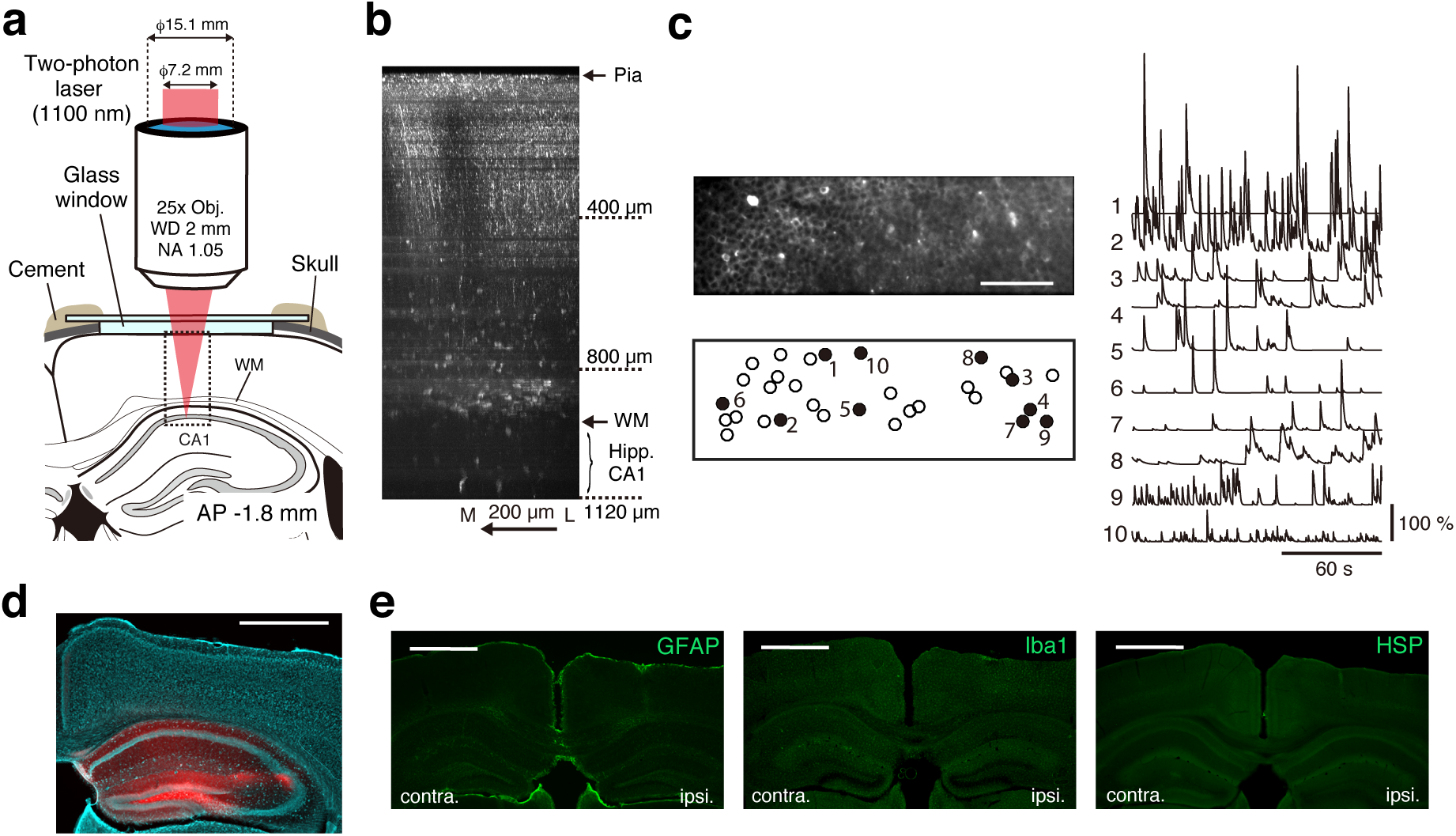
Optical access to hippocampal CA1 region in the intact brain. (**a**) Schematic illustration of *in vivo* two-photon imaging of the hippocampal CA1 region. The dotted square indicates the location where Z-stack images, as in (**b**), were acquired. (**b**) Representative XZ image of jRGECO1a-expressing hippocampus and deep cortical layer. Scale bar, 200 μm. (**c**) Left, top: representative time-averaged XY image of jRGECO1a-expressing CA1 pyramidal layer at a depth of 1040 μm from the cortical surface. Scale bar, 100 μm. Dense distribution of CA1 neurons is apparent when the images are time-averaged. Left, bottom: spatial distribution of identified neurons. Traces of the calcium transients of the numbered filled circles are shown on the right. (**d**) Expression of jRGECO1a and cell nuclei after 40 min (in total) imaging of the hippocampal CA1 region for 2 days. Scale bar, 1 mm. (**e**) Expressions of GFAP (left), Iba1 (middle), or HSP70/72 (right) in the contra-lateral and imaged hemispheres from the same mouse imaged in (**d**). Scale bar, 1 mm.

To demonstrate the utility of this method for identifying neural functions in deep areas in the intact brain, we examined reward prediction-related activity in the mFrC over ∼1 mm depth, as the mFrC demonstrates strong activity before movement starts (Friedman et al., 2015; Kim et al., 2016; Pinto and Dan, 2015; Sul et al., 2011). Head-restrained mice were conditioned to the delivery of a drop of water with an inter-delivery interval of 20 s (Figure 3a). As each session progressed (one session per day), the lick response to the water delivery increased in rate and became faster (Figure 3a–d). From the fifth session onwards, we performed two-photon calcium imaging of the mFrC at cortical depths of 100–1200 μm (Video 3). The imaging fields were classified into three areas according to depth (Paxinos and Franklin, 2007): the superficial area (100–300 μm, corresponding to layer 2/3 in the secondary motor area, M2), middle area (300–800 μm, corresponding to layer 5 in M2), and deep area (800–1200 μm, roughly corresponding to layer 6 in M2 and the PL area). In all three areas, approximately 50% of neurons showed a peak of mean (trial-averaged) activity during 5 s after the water delivery (Figure 3e, f), which was presumably related to licking and water acquisition (Figure 3b). Additionally, approximately 30% of neurons in all three areas showed a peak mean activity during 10 s before the water delivery (pre-reward period). The sequential distribution of the times of peak activity was not an artifact of ordering the neurons according to the time of peak activity, as the ratio of the mean activity around the peak activity to the baseline activity (ridge-to-background ratio; Harvey et al., 2012) was significantly higher than that of shuffled data (Figure 3g, h). Additionally, the sequential distribution of neurons with peak activity during the pre-reward period was not an artifact (Figure 3–Figure supplement 1). Even when 5 s windows were chosen from the pre-reward period, the ridge-to-background ratios of deep-area neurons with peak activity during each 5 s window were frequently higher than those in the shuffled data (Figure 3–Figure supplement 2 and Table supplement 1). To determine whether the activity pattern across trials was stable for individual neurons with peak activity during the pre-reward period, we calculated the correlation coefficient between the times of peak activity of two randomly separated groups of trials (Figure 3–Figure supplement 3a, b; see details in Materials and Methods section) and found that it was higher in the deep area than in the superficial area (Figure 3–Figure supplement 3c). This suggests that the PL neurons reliably code the reward prediction (or prediction of timing of the water delivery). The deep imaging method should therefore help us to understand the hierarchical and/or parallel processing occurring across the M2 and PL areas during decision-making.

**Figure 3.**
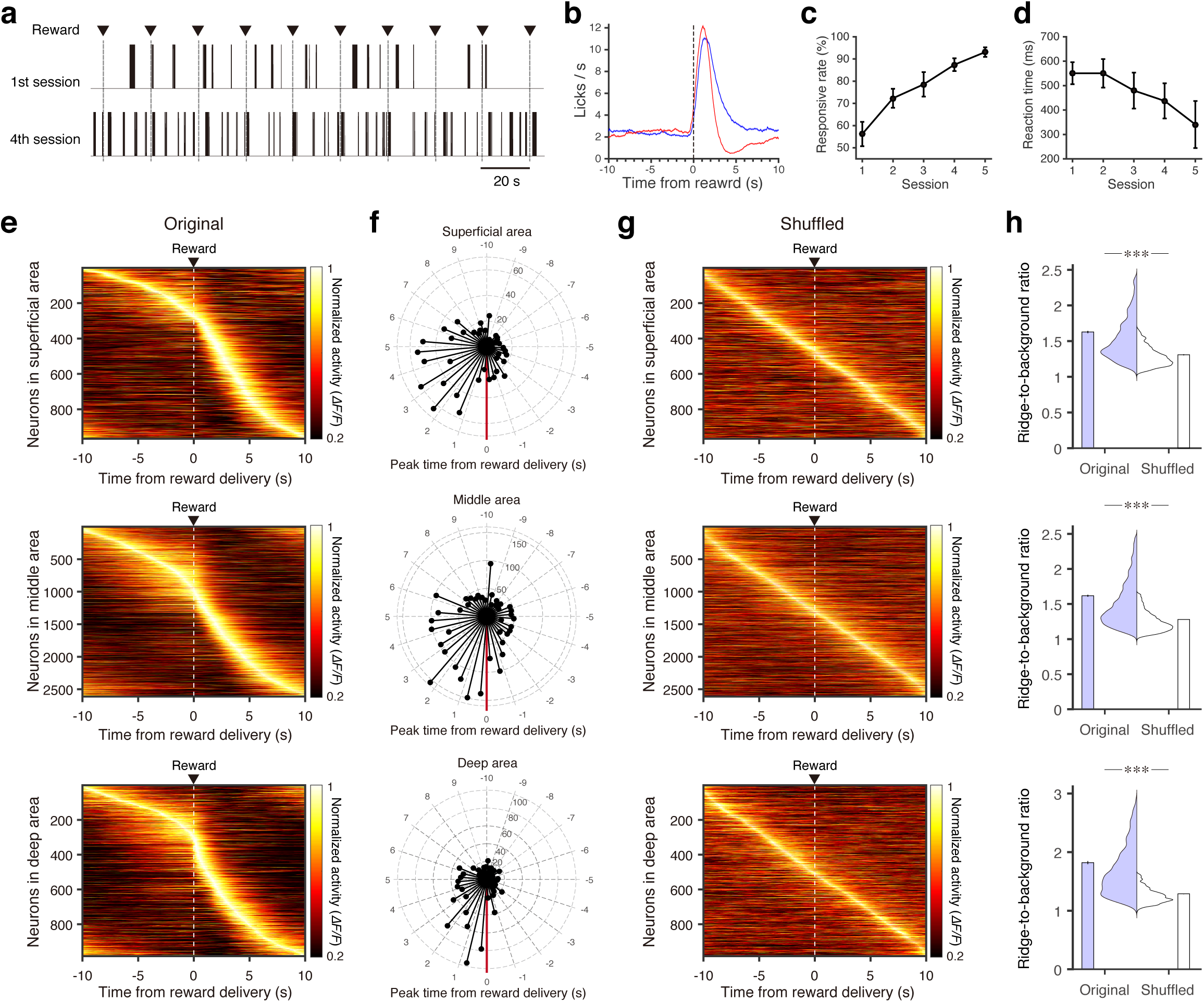
Neural activity in the medial frontal cortex during conditioning. (**a**) Representative traces of licking behavior at conditioning sessions 1 (top) and 4 (bottom) from the same mouse. Dashed-vertical lines indicate timings of water delivery. (**b**) Mean traces of lick frequency at sessions 1 (red) and 4 (blue; *n* = 11 mice). Light shading indicates the s.e.m. (**c**) Time course of responsive rate (rate of those deliveries with licking occurring within 2 s after the delivery; *n* = 11 at sessions 1–4 and *n* = 5 at session 5 before imaging started). (**d**) Time course of reaction time (time from the water delivery to the first lick). (**e**) Normalized trial-averaged activity of each neuron aligned with the water delivery (dashed lines) and ordered by the time of peak activity. Top, middle, and bottom panels are the superficial (12 fields from 7 mice), middle (35 fields from 10 mice), and deep (15 fields from 7 mice) areas respectively. (**f**) Polar histograms of the time of peak activity from reward delivery (red line). The time bin is 0.5 s, and they are ordered clockwise from the top (-10 s to 10 s). (**g**) Normalized trial-averaged shuffled activity of each neuron. The shuffled activity was calculated by circular shifts of the original calcium traces in each trial. (**h**) Distribution and mean of the ridge-to-background ratios in original and shuffled datasets. Top to bottom rows correspond to the superficial (*P* = 4.81 × 10^−101^, *n* = 966 neurons, Wilcoxon rank-sum test), middle (*P* = 4.34 × 10^−248^, *n* = 2612 neurons), and deep areas (*P* = 2.04 × 10^−146^, *n* = 983 neurons). ***: *P* < 0.001.

We also examined the activity pattern of hippocampal CA1 neurons in the young mice during the conditioning session (Video 4). The distribution of the time of peak activity was not random in all neurons, but was random in those neurons with a peak activity during the pre-reward period (Figure 4a–d). The time of peak activity of the latter neurons was also unstable across trials (Figure 4e). This indicates that the hippocampal CA1 neurons do not code the reward prediction in this conditioning without environmental changes, although we could not exclude the possibility that reward prediction-related activity may mature in adulthood.

**Figure 4.**
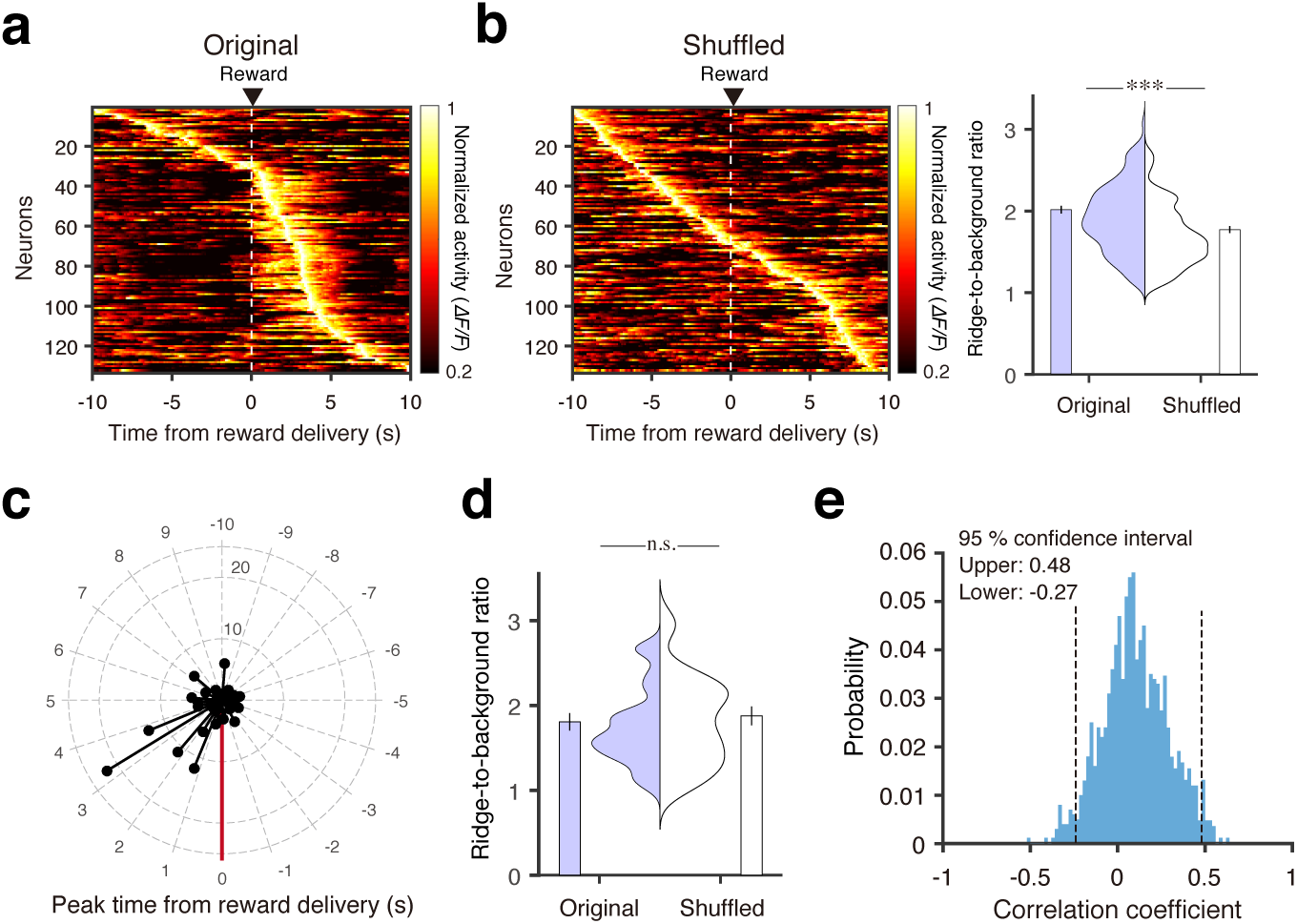
Neural activity in the hippocampal CA1 region during conditioning. (**a**) Normalized trial-averaged activity of each neuron aligned with the water delivery (dashed lines) and ordered by the time of peak activity. (**b**) Left, Normalized trial-averaged shuffled activity of each neuron aligned with the water delivery timing and ordered by the time of peak activity. Right, Distribution and mean of the ridge-to-background ratio in the original and shuffled datasets. ***: *P* = 0.000033, *n* = 133 neurons, Wilcoxon rank-sum test. (**c**) Polar histogram of the time of peak activity from reward delivery in each cell in the pooled data. (**d**) Distribution and mean of the ridge-to-background ratio of neurons with peak activity during the pre-reward period in original and shuffled datasets. *P* = 0.28, *n* = 31 neurons, Wilcoxon rank-sum test. (**e**) Histograms of the correlation coefficients of the time of peak activity between the two randomly divided groups of trials in neurons with peak activity during the pre-reward period.

In the intact brain, it is easy to change the field of view horizontally to the cortical surface. An 8 mm-wide glass window can be used for long-term imaging of the whole dorsal neocortex in the mouse (Kim et al., 2016), which would make it possible to image the medial prefrontal cortex and the hippocampus in memory-guided decision-making. If an objective with a wide field of view (>3 mm) is used (Sofroniew et al., 2016; Stirman et al., 2016; Tsai et al., 2015), the neural activity in both areas could be simultaneously imaged. Deep and wide-field two-photon calcium imaging of the intact brain will substantially help us understand the brain circuits that underlie cognition and memory processes.

## Competing interests

The authors declare no competing financial interests.

## Author contributions

M.K. and M.M. designed the experiments. M.K. performed two-photon imaging and mouse experiments and analyzed the data. M.O. and J. N. created R-CaMP1.07. K.K. made AAV1-hSyn-R-CaMP1.07. M.K. and M.M. wrote the paper, with comments from all the authors.

## Acknowledgements

We thank Y. Hirayama for technical assistance, J. Saito, Y.H. Tanaka, and H. Wake for plasmid construction, and Y.R. Tanaka for helpful discussion and writing a Matlab function to directly deal with images in the Olympus format. We thank V. Jayaraman, D.S. Kim, L.L. Looger, and K. Svoboda from the GENIE Project, Janelia Research Campus, Howard Hughes Medical Institute for providing AAV1-hSyn-jRGECO1a. This work was supported by Grant-in-Aids for Scientific Research on Innovative Areas (15H01455 to M.M.) and for Scientific Research (A) (15H02350 to M.M.) from the Ministry of Education, Culture, Sports, Science, and Technology, Japan, the Strategic Research Program for Brain Sciences and the program for Brain Mapping by Integrated Neurotechnologies for Disease Studies (Brain/MINDS) from Japan Agency for Medical Research and Development to M.M., and a Takeda Foundation grant to M.M.

## Materials and Methods

### Animals

All animal experiments were approved by the Institutional Animal Care and Use Committee of The University of Tokyo, Japan. All mice were provided with food and water *ad libitum* and housed in a 12:12 h light–dark cycle. The mice were not used for other experiments before this study. Male C57BL/6 mice (aged 2–3 months, SLC, Shizuoka, Japan) were utilized for mFrC imaging. Male and female C57BL/6 mice (aged 12–40 days in the young mice, and 2–3 months in the adult mice; Japan SLC, Shizuoka, Japan) were utilized for the imaging experiments on the hippocampus. For experiments using young mice, pups were weaned at P30, and then group-housed until the imaging window was implanted.

### Virus production

In this study, two red-fluorescent genetically encoded calcium indicators, R-CaMP1.07 (Ohkura et al., 2012) and jRGECO1a (Dana et al., 2016), were used. For imaging of R-CaMP1.07, the GCaMP3 DNA of pAAV-human synapsin I promoter (hSyn)-GCaMP3-WPRE-hGH polyA (Masamizu et al., 2014) was replaced with R-CaMP1.07 DNA from a pN1-R-CaMP1.07 vector construct (Ohkura et al., 2012). rAAV2/1-hSyn-R-CaMP1.07 (1.3 × 10^13^ vector genomes/ml) was produced with pAAV2-1 and purified as described previously (Kaneda et al., 2011; Kobayashi et al., 2016). rAAV2/1-hSyn-NES-jRGECO1a (2.95 × 10^13^ vector genomes/ml) was obtained from the University of Pennsylvania Gene Therapy Program Vector Core.

### Surgical procedures

#### mFrC

Mice were anesthetized by intramuscular injection of ketamine (74 mg/kg) and xylazine (10 mg/kg) before an incision was made in the skin covering the neocortex. After the mice were anesthetized, atropine (0.5 mg/kg) was injected to reduce bronchial secretion and improve breathing, and an eye ointment (Tarivid; 0.3% w/v ofloxacin, Santen Pharmaceutical, Osaka, Japan) was applied to prevent eye-drying. Body temperature was maintained at 36–37°C with a heating pad. After the exposed skull was cleaned, a head-plate (Tsukasa Giken, Shizuoka, Japan; Hira et al., 2013) was attached to the skull using dental cement (Fuji lute BC; GC, Tokyo, Japan, and Bistite II; Tokuyama Dental, Tokyo, Japan). The surface of the intact skull was coated with dental adhesive resin cement (Super bond; Sun Medical, Shiga, Japan) to prevent drying. An isotonic saline solution with 5 w/v% glucose was injected intraperitoneally after the surgery. Mice were allowed to recover for 1 day before virus injection.

Thirty minutes before surgery for virus injection, dexamethasone sodium phosphate (1.32 mg/kg) was administered intraperitoneally to prevent cerebral edema. Mice were anesthetized with isoflurane (3–4% for induction and ∼1% during surgery) inhalation and placed on a stereotaxic frame (SR-5M; Narishige, Tokyo, Japan). Before virus injection, a pulled glass pipette (broken and beveled to an outer diameter of 25–30 μm; Sutter Instruments, CA, USA) and a 5 μl Hamilton syringe were back-filled with mineral oil (Nacalai Tesque, Kyoto, Japan) and front-loaded with virus solution. The virus solution was then injected into the mFrC (2.7–2.8 mm anterior and 0.4 mm left of the bregma, 800–1200 μm dorsal from the pial surface). To minimize background fluorescence from solution backflow through the space made by the glass capillary insertion, the axis of the glass capillary was angled 30–40° from the horizontal plane. From 100 to 200 nl of AAV solution was injected via a syringe pump at a rate of 15–20 nl/min (KDS310; KD Scientific, MA, USA). The capillary was maintained in place for more than 10 min after the injection before being slowly withdrawn. The craniotomy was then covered with silicon elastomer (quick cast, World Precision Instruments, FL, USA) and dental adhesive (Super bond). At least 3 weeks after the viral injection, the craniotomy (1.5 mm diameter at the area of interest) was conducted and dura mater was removed. A glass window was placed over the craniotomy, and the edge was sealed with cyanoacrylate adhesive (Vetbond, 3M, MN, USA), dental resin cement, and dental adhesive. The glass window consisted of two circular cover slips (No.1, 0.12–0.17 mm thickness and 2.5 mm diameter; and No.5, 0.45–0.60 mm thickness and 1.5 mm diameter; Matsunami Glass, Osaka, Japan) that were glued together with UV-curing optical adhesive (NOR-61; Norland Products, NJ, USA). After the window implantation, a 250 μl saline solution containing anti-inflammatory and analgesic carprofen (6 mg/kg) was administered intraperitoneally. Mice were then returned to their cages, and imaging sessions were started after allowing at least 1 day for recovery.

#### Hippocampus

The procedures for the hippocampus were mostly the same as those for the mFrC. However, when the viral solution was injected at P12–14, the head-plate was not attached, as at their age the body size was too small to allow attachment. Two weeks after injection, the mice were anesthetized by intraperitoneal injection of ketamine (74 mg/kg) and xylazine (10 mg/kg), the head-plate was attached, and the imaging window was implanted. The glass window consisted of two circular cover slips (No.1, 0.12–0.17 mm thickness and 2.5 mm diameter; and No.3, 0.25–0.35 mm thickness and 1.5 mm diameter). The dura mater was not removed, as it was thinner and more fragile than that in the adult mice.

## Behavioral conditioning

The mice were water-deprived in their home cages and maintained at 80–85% of their normal weight throughout the experiments. During the behavioral conditioning, mice were set within a body chamber and head-fixed with custom-designed apparatus (O’Hara, Tokyo, Japan; Hira et al., 2013). A spout was set in front of their mouth, and a 4 μl drop of water was delivered from the spout at a time interval of 20 s. The mice were allowed to lick at any time, and licking behavior was monitored by an infrared LED sensor. The rate of water delivery that incurred at least one lick during 2 s after the delivery was defined as the responsive rate. The duration of the daily conditioning sessions was 40–60 min. At the end of each session, the mice were allowed to freely gain water drops (total water consumption was ∼1 ml per session). On rest days (typically weekends), the mice had free access to a 3% agarose block (1.2 g per day) in the cage.

## Two-photon calcium imaging

Two-photon imaging was conducted using an FVMPE-RS system (Olympus, Tokyo, Japan) equipped with a 25× water immersion objective (for imaging of the mFrC: XLPLN25XSVMP, numerical aperture: 1.0, working distance: 4 mm, Olympus; for imaging of the hippocampus: XLPLN23XWMP2, numerical aperture: 1.05, working distance: 2 mm, Olympus) and a broadly tunable laser with a pulse width of 120 fs and a repetition rate of 80 MHz (Insight DS+ Dual, Spectra Physics, CA, USA), set at a wavelength of 1100 nm. Fluorescence emissions were collected using a GaAsP photomultiplier tube (Hamamatsu Photonics, Shizuoka, Japan). To shorten the light-path length within the tissue, the back aperture of the objective was underfilled with the diameter-shortened (7.2 mm, in comparison with that of the back aperture of 15.1 mm or 14.4 mm) laser beam^11^. When the modified laser at a wavelength of 1100 nm was used for two-photon excitation of 0.1 μm fluorescent beads, the full-widths at half-maximum were 0.94 ± 0.09 μm (mean ± s.d., *n* = 5 beads) laterally and 7.7 ± 1.70 μm (*n* = 5 beads) axially, which are comparable to those used for two-photon calcium imaging of multiple neurons with cellular resolution (Lecoq et al., 2014; Sadakane et al., 2015).

During the imaging session, the mouse head was fixed and the body was constrained within a body chamber under the microscope (OPR-GST, O’Hara; Masamizu et al., 2014). Before the first imaging session of each mouse started, the angle of the stage on which the mouse chamber was placed was finely adjusted to set the glass window perpendicular to the optical axis. This was accomplished by the imaging of microbeads on the surface of the glass window (Kawakami et al., 2015). The frame acquisition rate was 30 frames/s, and the size of the imaging fields was generally 512 × 512 pixels (0.904 μm/pixel), or 512 × 160 pixels, with three-frame averaging to increase the signal-to-noise ratio. The depth of the functional imaging plane was up to 1200 μm from the cortical surface (*n* = 62 planes in the mFrC from 11 mice expressing R-CaMP1.07, *n* = 6 in the hippocampus from three mice expressing jRGECO1a). The duration of one imaging session was 15–20 min, and 1–4 imaging sessions from different depths were performed in a daily experiment. For each mouse, imaging was conducted for 3–5 days.

## Image processing

Analyses were performed using MATLAB (R2016a, version 9.0.0.341360; MathWorks, MA, USA) and Fiji software (Schindelin et al., 2012). Raw image sequences acquired on the FVMPE-RS system were loaded into MATLAB using custom-written scripts. Motion correction was performed by phase-correlation using the Suite2P package (Pachitariu et al., 2016). After the motion correction, images were three frame-averaged before being analyzed. A constrained non-negative matrix factorization (cNMF) algorithm was employed to extract neural activities from a time series of images (Pnevmatikakis et al., 2016). The noise variances in the power spectrum density at high frequency estimated by the cNMF algorithm were as follows (mean ± s.d.): 14.42 ± 5.11 (*n* = 12 fields) in the superficial area of the mFrC, 22.00 ± 10.27 (*n* = 35 fields) in the middle area of the mFrC, 21.69 ± 12.63 (*n* = 15 fields) in the deep area of the mFrC, and 19.45 ± 3.68 (*n* = 6 fields) in the hippocampal CA1 region. The detrended relative fluorescence changes (Δ*F*/*F*) were calculated with eight percentile values over an interval of ±30 s around each sample time point (Dombeck et al., 2007). Traces of Δ*F*/*F* from 10 s before to 10 s after the water delivery in those deliveries with at least one lick during 2 s after the delivery were used for the analyses.

## Data analysis

The ridge-to-background ratio was used for the estimation of the distribution of the time of peak activity (Harvey et al., 2012). To create a shuffled Δ*F*/*F* trace of each neuron, the time point of the actual Δ*F*/*F* trace was circularly shifted by a random amount for each trial and then trial-averaged. For each neuron, the ridge Δ*F*/*F* was defined as the mean Δ*F*/*F* over 12 frames (100 ms/frame) surrounding the time of peak activity, and the background Δ*F*/*F* was defined as the mean Δ*F*/*F* in the other data points. The ridge Δ*F*/*F* was then divided by the background Δ*F*/*F*.

The trial-by-trial stability of the time of peak activity of the neurons that had their peak activity during the pre-reward period (-10 s to 0 s) was evaluated as follows: in each session, all trials were randomly divided into two groups, and the trial-averaged activity in each group was calculated for each neuron. To remove the effects of different sample sizes across the three mFrC areas and the hippocampus, 50 neurons were randomly chosen from all imaging fields in each area. The time of peak activity in one group was plotted against that in the other, and the Pearson’s correlation coefficient was determined. Thus, if the timing of the peak activity of each neuron was constant across trials, the correlation coefficient should be 1. This procedure was repeated 1000 times, and the 95% confidence interval was determined for each of the areas. When the lower bound of the 95% confidence interval was above zero, it was concluded that the time of peak activity was not random across trials. To estimate the difference in the trial-by-trial stability of the time of peak activity between pairs of the three areas in the mFrC (Figure 3–Figure supplement 3c), the mean correlation coefficients were compared using a permutation test. For each pair from the superficial, middle, and deep areas, all neurons with peak activity during the pre-reward period were randomly reassigned to one of the two areas. For each area with reassigned neurons, the correlation coefficient between the times of peak activity of the two randomly separated groups of trials was calculated, and the absolute difference of the correlation coefficients between the two areas was estimated. This procedure was repeated 10000 times, and the distribution of the absolute differences between the two areas was determined. Following this, the statistical significance was determined according to whether or not the absolute difference in the mean correlation coefficients between the two areas with original neurons assigned (Figure 3–Figure supplement 3b) was above the 95th percentile of the resampled distribution corrected using the Bonferroni method.

## Histology

After the last *in vivo* imaging session, the mice were deeply anesthetized with ketamine (74 mg/kg) and xylazine (10 mg/kg) and transcardially perfused with 40 ml of phosphate buffered saline (PBS) and 40 ml of 4% paraformaldehyde in PBS (Wako, Osaka, Japan). The brains were removed and postfixed with the same fixative at 4°C for longer than 12 h. For immunostaining, the brains were cut into coronal sections with a thickness of 50–100 μm.

Slices were washed in PBS-X (0.5% triton-X in PBS) containing 10% normal goat serum, and then incubated with one of the primary antibodies (1:500 dilution of rabbit anti-GFAP [glial fibrillary acidic protein], G9269, Sigma-Aldrich, MO, USA; 1:500 dilution of rabbit anti-Iba1, 019-19741, Wako; 1:400 dilution of mouse anti-HSP70/72 [heat shock protein 70/72], ADI-SPA-810-F, Enzo Life Sciences, NY, USA) overnight at 4°C. Afterwards, slices were washed in PBS-X and incubated with species-appropriate Alexa Fluoro-488 conjugated secondary antibody (1:500 dilution of anti-rabbit IgG for GFAP and Iba1 antibodies; 1:500 dilution of anti-mouse IgG for HSP70/72 antibody). After staining the cell nuclei with fluorescent Nissl stain (NeuroTrace 435/455, 1:200, N21479, Thermo Fisher Scientific, MA, USA), the slices were mounted on glass slides with Fluoromount/Plus mounting medium (Diagnostic BioSystems, CA, USA). Fluorescence images were acquired with an upright fluorescence microscope (BX53, Olympus) and a CCD camera (Retiga 2000R, Q Imaging, BC, Canada), and analyzed with Fiji software (Schindelin et al., 2012).

## Statistics

Data are presented as mean ± s.d., and the Wilcoxon rank-sum tests, Spearman’s correlation tests, Pearson’s correlation tests, and permutation tests described above were used for statistical comparisons. Pairwise comparisons were two-tailed unless otherwise noted. Error bars in graphs represent the s.e.m. No statistical tests were run to predetermine the sample size, and blinding and randomization were not performed.

---

**Video 1. | Representative two-photon XYZ images of the mFrC expressing R-CaMP1.07.**

The depth increment in the image stack was 2.0 μm, and the bottom depth of the imaging was 1100 μm. The field of view was 512 × 512 pixels with a size of 509.18 μm × 509.18 μm. Each image was the average of 16 frames. The mouse was not anesthetized. The motion correction was not conducted. The movie was denoised with a spatial Gaussian filter (σ = 0.6). The right image corresponds to XZ plane of the XYZ images (max intensity-projection toward Y dimension) and the horizontal yellow line indicates the current depth of the left XY image.

**Video 2. | Representative two-photon XYZ images of the neocortex and hippocampus that expressed jRGECO1a.**

The depth increment in the image stack was 2.5 μm, and the bottom depth of the imaging was 1100 μm. The field of view was the same size as in Video 1. Each image was the average of 16 frames. The mouse was not anesthetized. The motion correction was not conducted. The movie was denoised with a spatial Gaussian filter (σ = 0.6). The right image corresponds to XZ plane of the XYZ images (max intensity-projection toward Y dimension) and the horizontal yellow line indicates the current depth of the left XY image. Some leakage of the virus from the hippocampus to the neocortex during the injection procedure may cause a subset of the neocortical neurons to express jRGECO1a.

**Video 3. | Functional imaging of the PL area expressing R-CaMP1.07 during conditioning.**

The imaging depth was 1100 μm from the cortical surface. The field of view was 509.18 μm × 509.18 μm and 512 × 512 pixels. White circles at the right bottom indicate the timing of water delivery, with an inter-delivery interval of 20 s. The data were acquired at 30 Hz, and the movie was downsampled to 5 Hz and denoised with a spatio-temporal Gaussian filter (spatial σ = 0.6, temporal σ = 0.8).

**Video 4. | Functional imaging of the hippocampal CA1 pyramidal layer that expressed jRGECO1a during conditioning.**

The imaging depth was 1000 μm from the cortical surface, and the upper cortical tissue was intact. The field of view was 509.18 μm × 159.04 μm and 512 × 160 pixels. White circles at the right bottom indicate the timing of water delivery, with an inter-delivery interval of 20 s. The data were acquired at 30 Hz (three frame-averaged), and the movie was downsampled to 5 Hz and denoised with a spatio-temporal Gaussian filter (spatial σ = 0.6, temporal σ = 0.8).

**Figure 3–Figure supplement 1.**
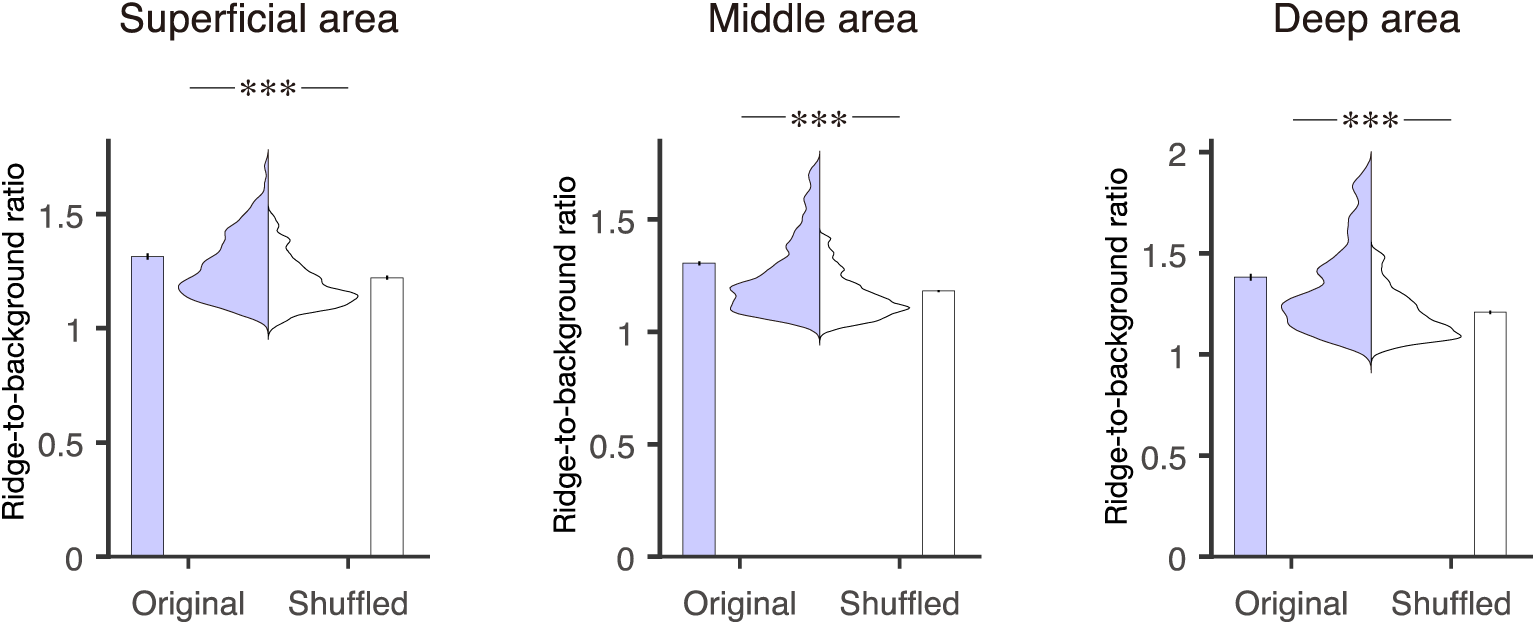
Ridge-to-background ratios of the mFrC neurons showing peak activity during the pre-reward period. Distribution and mean of the ridge-to-background ratios in original and shuffled datasets from neurons with peak activity during the pre-reward period. Left to right panels correspond to the superficial (*P* = 1.99 × 10^−8^, *n* = 273 neurons, Wilcoxon rank-sum test), middle (*P* = 4.21 × 10^−37^, *n* = 943 neurons), and deep areas (*P* = 2.37 × 10^−17^, *n* = 286 neurons). ***: *P* < 0.001.

**Figure 3–Figure supplement 2.**
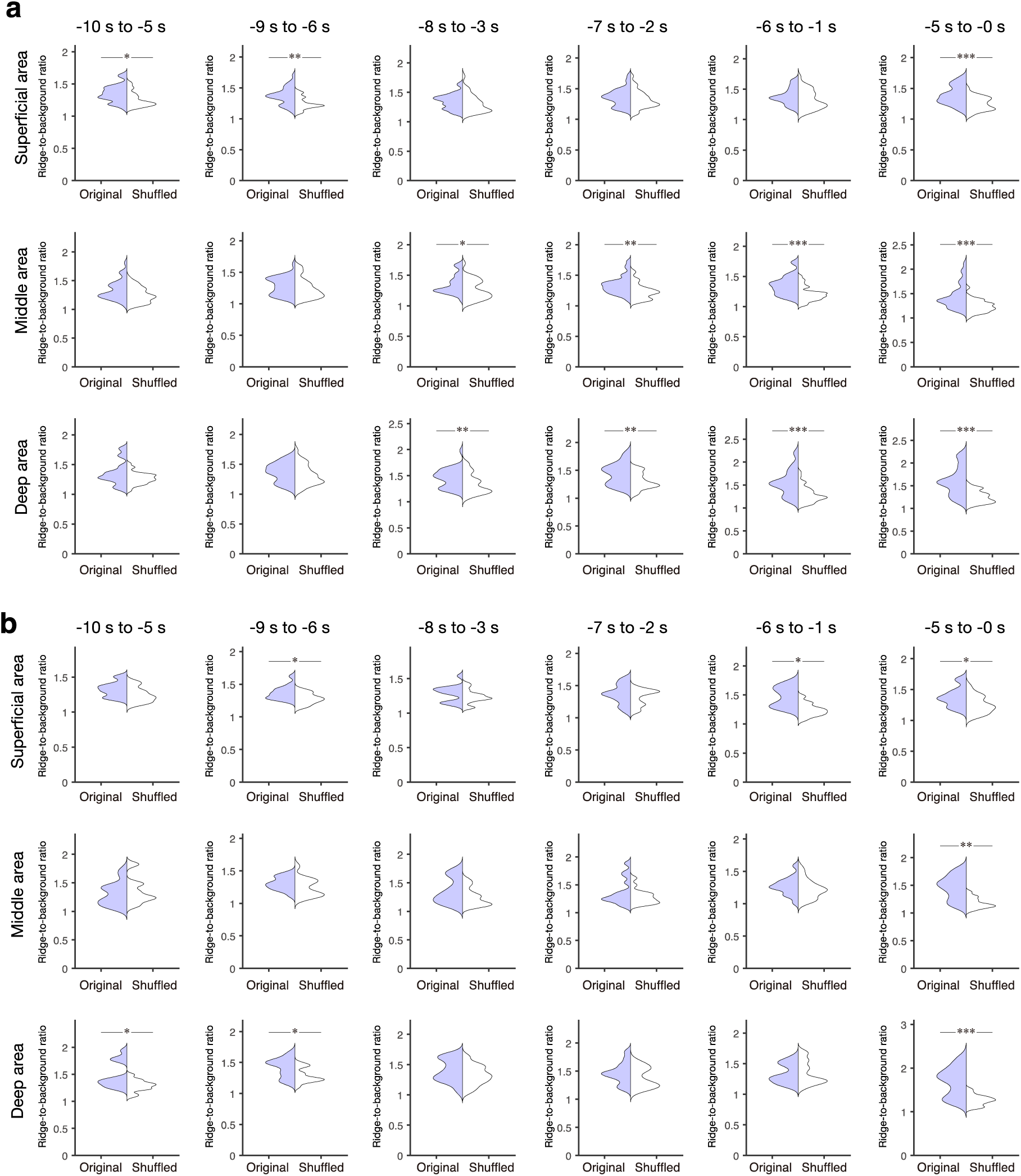
Ridge-to-background ratios of neurons showing peak activity during 5 s windows in the pre-reward period. (**a, b**) Examples of ridge-to-background ratios calculated from neurons with peak mean activity occurring within six 5 s time windows (from left to right: −10 s to −5 s, −9 s to −4 s, −8 s to −3 s, −7 s to −2 s, −6 s to −1 s, and −5 s to 0 s). Thirty-five (**a**) and fifteen (**b**) neurons were randomly chosen to calculate the ratios. Top, middle, and bottom panels are the superficial, middle, and deep areas of the mFrC, respectively. *: *P* < 0.05, **: *P* < 0.01, ***: *P* < 0.001, Wilcoxon rank-sum test.

**Figure 3–Figure supplement 3.**
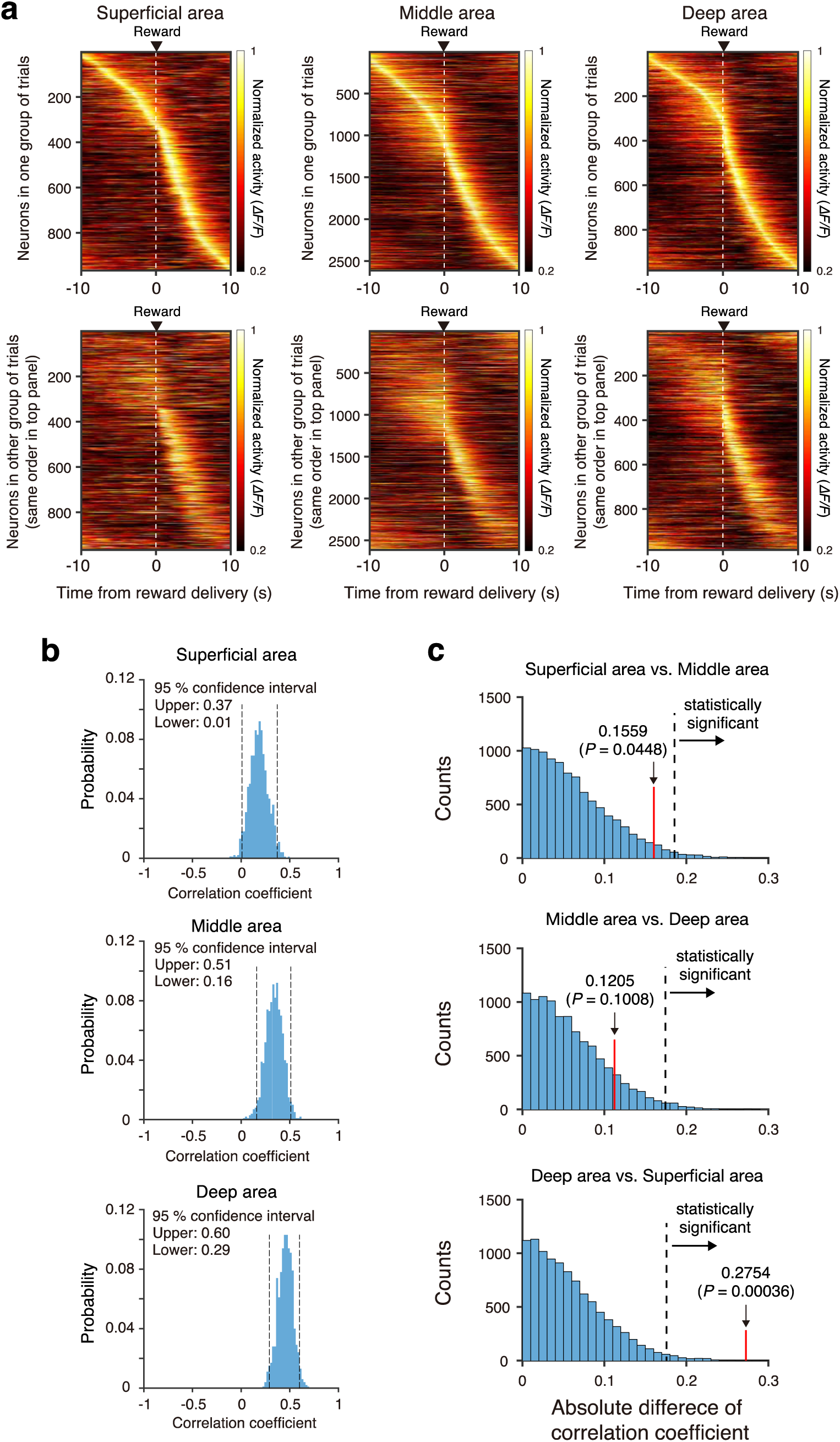
Stability of trial-by-trial activity of the mFrC neurons with peak activity during the pre-reward period. (**a**) Mean activity of mFrC neurons around the water delivery (dashed lines) from two representative separated groups of trials (top and bottom). In each column, neurons were ordered by the time of peak activity in the top group. A high similarity in the distribution between the top and bottom panels implies that trial-by-trial activity is stable. Left, the superficial area. Middle, the middle area. Right, the deep area. (**b**) Histograms of correlation coefficients of the time of peak activity between two randomly divided groups of trials, for those neurons with peak activity during the pre-reward period (−10 s to 0 s). The random division of trials was performed 1000 times (see details in Materials and Methods section). (**c**) Distribution of the absolute difference of the correlation coefficient between two areas (top, superficial area vs. middle area; middle, middle area vs. deep area; bottom, deep area vs. superficial area) with randomly reassigned neurons with peak activity during the pre-reward period. The difference was estimated by a permutation procedure (see details in Data Analysis section in Materials and Methods). Dashed lines indicate 95th percentile corrected by the Bonferroni method (resulting in a 98.3% position). Red lines indicated the absolute differences of mean correlation coefficients calculated from (**b**). In the bottom subimage, the red line is right of the dashed line, indicating that the trial-to-trial stability was statistically higher in the deep area than in the superficial area.

**Figure 3–Table supplement 1.**
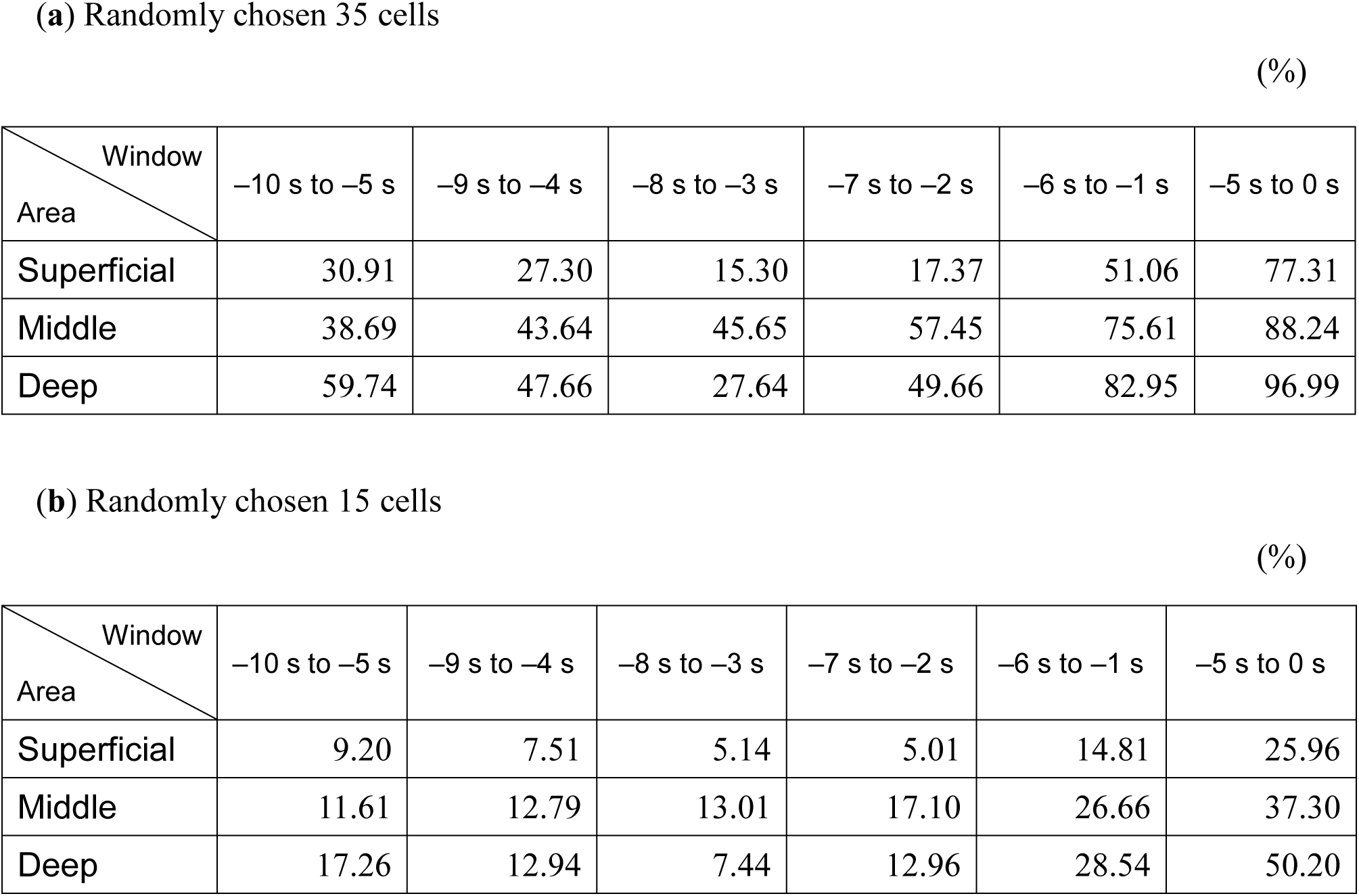
Statistics of ridge-to-background ratios of neurons showing peak activity during the 5 s windows in the pre-reward period. Percentages of ridge-to-background ratios in which original data were significantly different (*P* < 0.05, Wilcoxon rank-sum test) from shuffled data. Random chooses of thirty-five (**a**) and fifteen (**b**) neurons to calculate the ratios were repeated 10000 times.

